# Non-autonomous Vnd acts with autonomous Appl during mushroom body axon growth in *Drosophila*

**DOI:** 10.1101/2023.09.01.555727

**Authors:** Claire Marquilly, Ana Boulanger, Germain U. Busto, Robert P Zinzen, Bassem A Hassan, Lee G Fradkin, Jean-Maurice Dura

## Abstract

The Amyloid Precursor Protein (APP) is associated with Alzheimer’s disease. *Appl* is the single *Drosophila* APP ortholog and is expressed in all neurons throughout development. Appl was previously shown to cell-autonomously modulate axon outgrowth in the mushroom bodies (MBs), the fly olfactory memory center. However, we found that *Appl*^*d*^, the only reported null allele, affects the normal function of *vnd*, the gene just proximal to *Appl*. To decipher developmental defects specifically due to a loss of only *Appl* function, we generated a precise *Appl* null allele (*Appl*^*C2*.*1*^) by CRISPR/Cas9 genomic engineering. With *Appl*^*C2*.*1*^, we confirmed the partial requirement for *Appl* in MB axon outgrowth. We also produced new CRISPR *vnd* alleles removing either *vnd-B* or *vnd-A* function. We report here that *vnd* is also required for MB β-branch axon outgrowth and to a much greater extent than *Appl* itself. Moreover, *vnd* is expressed in neurons close to, but not within, the MB during development and is required non-cell-autonomously for MB axon outgrowth

## Introduction

Alzheimer’s disease (AD) is associated with extracellular accumulation of amyloid fibrils derived from the Amyloid Precursor Protein (APP). APPs have therefore been intensely investigated, however their physiological function in the brain remains unclear and controversial (Nicolas and Hassan, 2014; Selkoe and Hardy, 2016; Soldano and Hassan, 2014). Although there are three paralogues in mammals (APP, APLP1 and APLP2), *Drosophila* encodes a single APP homologue, called *Appl*, that is expressed in all neurons throughout development. It has been shown that the type I transmembrane protein coding gene *Appl* is a conserved neuronal modulator of a Wnt planar cell polarity (Wnt/PCP) pathway, a regulator of cellular orientation within the plane of an epithelium (Soldano et al., 2013). It has been proposed that Appl is part of the membrane complex formed by the core PCP receptors Fz (*frizzled*) and Vang (*Van Gogh*) required for axon growth (Liu et al., 2021; Soldano et al., 2013).

The mushroom bodies (MBs) are two bilaterally symmetric structures in the central brain that are required for learning and memory (Lin, 2023). Each MB is comprised of 2000 neurons that arise from 4 neuroblasts. Three types of neurons appear sequentially during development: the embryonic/early larval γ, the larval α’β’, and the late larval/pupal αβ types. Each αβ neuron projects an axon that branches to send an α branch dorsally, which contributes to the formation of the α lobe, and a β branch medially, which contributes to the formation of the β lobe (Lee et al., 1999). Appl’s expression continues in adult flies, notably in the MB αβ neurons (Leyssen et al., 2005; Rabah et al., 2025). So far, *Appl*^*d*^ is the only reported null *Appl* allele and it results from a synthetic genomic deletion removing the *Appl* locus (Luo et al., 1992). The *Appl*^*d*^-bearing chromosome was selected, after γ irradiation, as a translocation of a partial duplication of the X chromosome on the Y chromosome to a X chromosome terminal deficiency (S1 Fig). *Appl*^*d*^ flies are viable, fertile and display no gross structural defects in the brain (Luo et al., 1992). However, the *Appl* signaling pathway is required for proper axon outgrowth in the MBs since *Appl*^*d*^ flies display modestly penetrant axonal defects in αβ neurons. Importantly, Appl is required cell-autonomously for β-branch axon outgrowth (Cassar and Kretzschmar, 2016; Soldano et al., 2013).

The *ventral nervous system defective* (*vnd*) gene, which is immediately proximal to *Appl*, encodes a Nk2-class homeodomain transcription factor, that acts in a context-dependent manner as an activator or repressor and is essential for the development of the nervous system. The *vnd* gene encodes two proteins: Vnd-A and Vnd-B whose mRNAs arise from two different promoters. These two proteins differ in their aminoterminal domains and are identical in the remainder of their sequences. While Vnd-A is a transcription repressor for promoters containing Nk-2 binding sites, Vnd-B directly activates transcription, also likely via binding to the same binding site. (Stepchenko et al., 2011). Flies bearing a null allele of vnd in hemi- or homozygous condition are embryonic lethal (Jimenez et al., 1995).

Interestingly, we found that the *Appl*^*d*^ chromosome also genetically affects *vnd* function. To genetically dissect *Appl* and *vnd* functions, we generated CRISPR alleles that precisely delete the *Appl* gene without affecting *vnd* function (*Appl*^*C2*.*1*)^ and that precisely delete each one of the two *vnd* transcripts without affecting *Appl* (*vnd*^*CΔA*^ and *vnd*^*CΔB*^). To analyze the exact developmental defects due to the specific loss of the *Appl* function only, we used *Appl*^*C2*.*1*^ and first confirmed the partial requirement of *Appl* in MB β-branch axon outgrowth. We showed here that *vnd-A*, but not *vnd-B*, is also required for MB β-branch axon outgrowth. Unexpectedly, *vnd* is expressed in neurons close to, but not within, the MB during development and is required non-cell-autonomously, likely by promoting the production of a secreted factor(s) involved in the MB axon outgrowth.

## Results

### Generating new *Appl* CRISPR alleles

The complex chromosome structure resulting in the *Appl*^*d*^ allele complicates its subsequent genetic manipulation and might be a reason for the sensitivity of its phenotype to genetic background (S1 Fig). Therefore, we produced new null *Appl* alleles by removing the entire *Appl* transcriptional unit via CRISPR/Cas9-mediated deletion (Doudna and Charpentier, 2014). We recovered *Appl*^*C1*.*4*^ and *Appl*^*C2*.*1*^, two CRISPR *Appl* null alleles (Fig 1 and S2 Fig), following an adaptation of a published protocol to produce ∼50kb deletions (Port et al., 2014). *Appl*^*d*^ MBs display a modestly penetrant cell-autonomous axon growth defect of the β-branches which manifests as an absence of the β lobe (Liu et al., 2021; Soldano et al., 2013).

**Fig 1.**
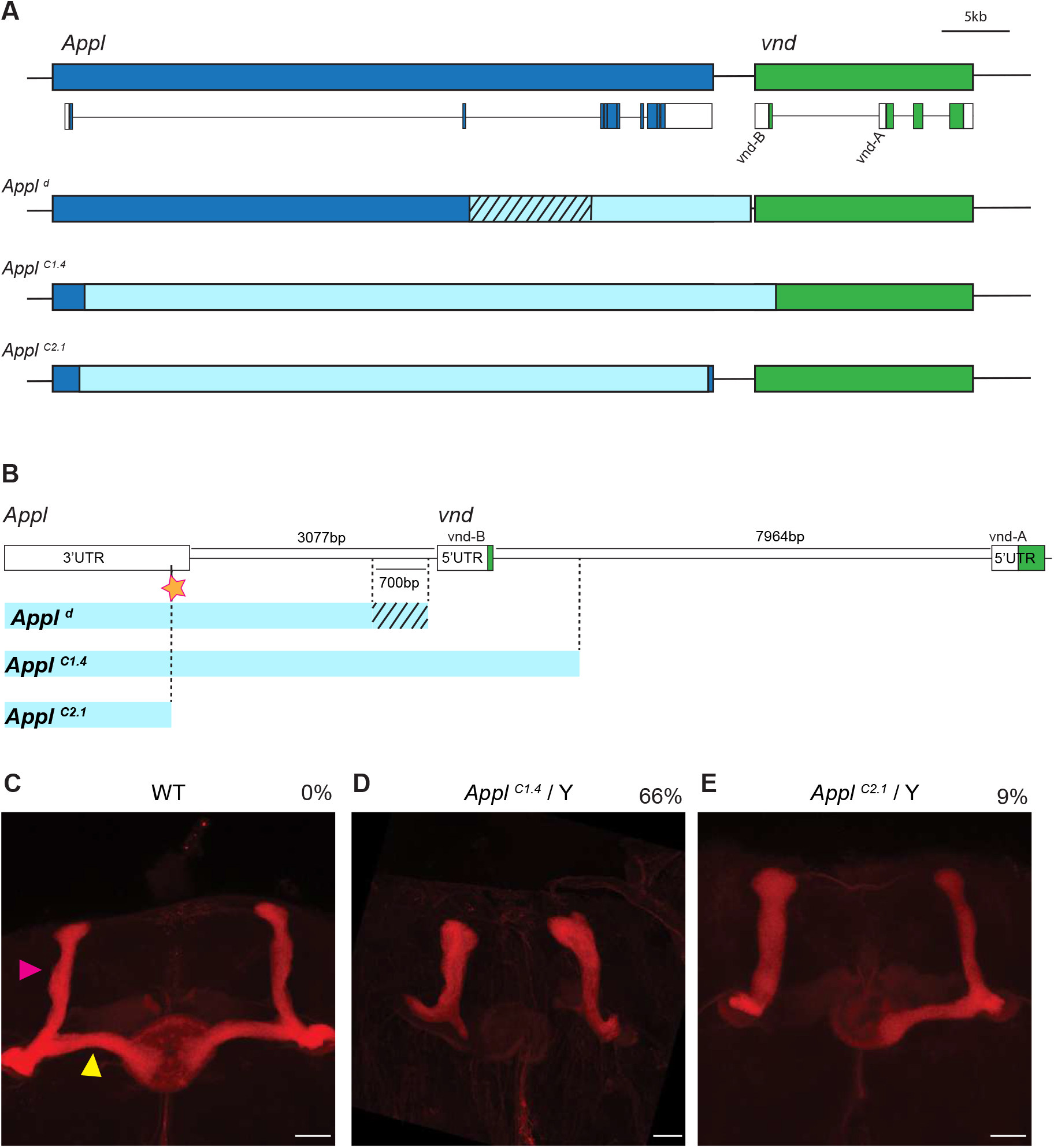
Generating new *Appl* CRISPR alleles. (A) Schematic representation of *Appl* (blue) and *vnd* (green) genes and transcripts. White boxes represent 5’UTR and 3’UTR, blue and green boxes represent *Appl* and *vnd* coding sequences respectively. Schematic representations of the *Appl*^*d*^, *Appl*^*C1*.*4*^ and *Appl*^*C2*.*1*^ mutant alleles where the deleted sequences are represented in light blue. The hatched black segment represents the uncertainty where the deletion of *Appl*^*d*^ starts at its 5’end. (B) Representation of the 3’ limit of the *Appl*^*d*^, *Appl*^*C1*.*4*^ and *Appl*^*C2*.*1*^ mutant alleles (light blue). The hatched black segment represents the uncertainty where the *Appl*^*d*^ deletion ends in 3’. The orange star represents the position of the 3’ sgRNA used to create the two CRISPR deletions. (C) Wild-type (WT) MB revealed by an anti-Fas2 staining. The pink and yellow arrowheads show respectively the α and β lobe (n=216 MBs). (D-E) Anti-Fas2 staining reveals the loss of β lobe on a representative *Appl*^*C1*.*4*^ brain (n=147 MBs) (D) and on *Appl*^*C2*.*1*^ brain (n=253 MBs) (E). The % represents the proportion of loss of β lobe for each genotype. Scale bar = 50 µm. Full genotypes: (C) Canton-S. (D) y *Appl*^*C1*.*4*^ *w*^*1118*^/Y. (E) *Appl*^*C2*.*1*^/Y.

Surprisingly, we found a strong difference in the penetrance of the absence of the β lobe phenotype in the two *Appl* CRISPR alleles. Although 9% of *Appl*^*C2*.*1*^ MBs lacked the β lobe, slightly lower than the 14.5% described for *Appl*^*d*^ (Marquilly et al., 2021), *Appl*^*C1*.*4*^ MBs displayed a much higher penetrance (66%) of this phenotype (Fig 1). We determined the extents of the precise deletions present in the two alleles via sequencing of PCR amplicons and found that while the *Appl*^*C2*.*1*^ deletion precisely removed the *Appl* transcriptional unit, the *Appl*^*C1*.*4*^ allele also removed a part of the *vnd* transcriptional unit (Fig 1). We then similarly mapped the *Appl*^*d*^ deletion and found that, unlike *Appl*^*C2*.*1*^, it removes most of the intergenic region between *Appl* and *vnd* which possibly influences *vnd* function (Fig 1). We then assessed, by genetic complementation tests, if the different *Appl* null alleles possibly affected *vnd* function. We used *vnd*^*A*^, a molecularly characterized lethal allele which impacts both the *vnd-A* and *vnd-B* transcripts (Fig 2) (Haelterman et al., 2014). Although *Appl*^*C2*.*1*^/*vnd*^*A*^ transheterozygous MBs displayed no anatomical MB phenotypes, the *Appl*^*d*^/*vnd*^*A*^ and *Appl*^*C1*.*4*^/*vnd*^*A*^ MBs displayed 14% and 69% of β lobe absence, respectively (Fig 2). Flies heterozygous for any of the four mutations individually displayed no MB phenotypes: *vnd*^*A*^/+ (n = 72), *Appl*^*C1*.*4*^/+ (n = 60), *Appl*^*C2*.*1*^/+ (n = 42) and *Appl*^*d*^/+ previously described (Soldano et al., 2013). Therefore, *Appl*^*d*^ and *Appl*^*C1*.*4*^, but not *Appl*^*C2*.*1*^ affect *vnd* functions in MB development. We could write *Appl*^*d*^ = *Appl*^*null*^ *vnd*^*weak*^, *Appl*^*C1*.*4*^ = *Appl*^*null*^ *vnd*^*strong*^ and only *Appl*^*C2*.*1*^ = *Appl*^*null*^ *vnd*^+^. Taken together, these data strongly suggest that *vnd* is also involved in the MB β-branch axon growth and likely to a much greater extent than *Appl* itself.

**Fig 2.**
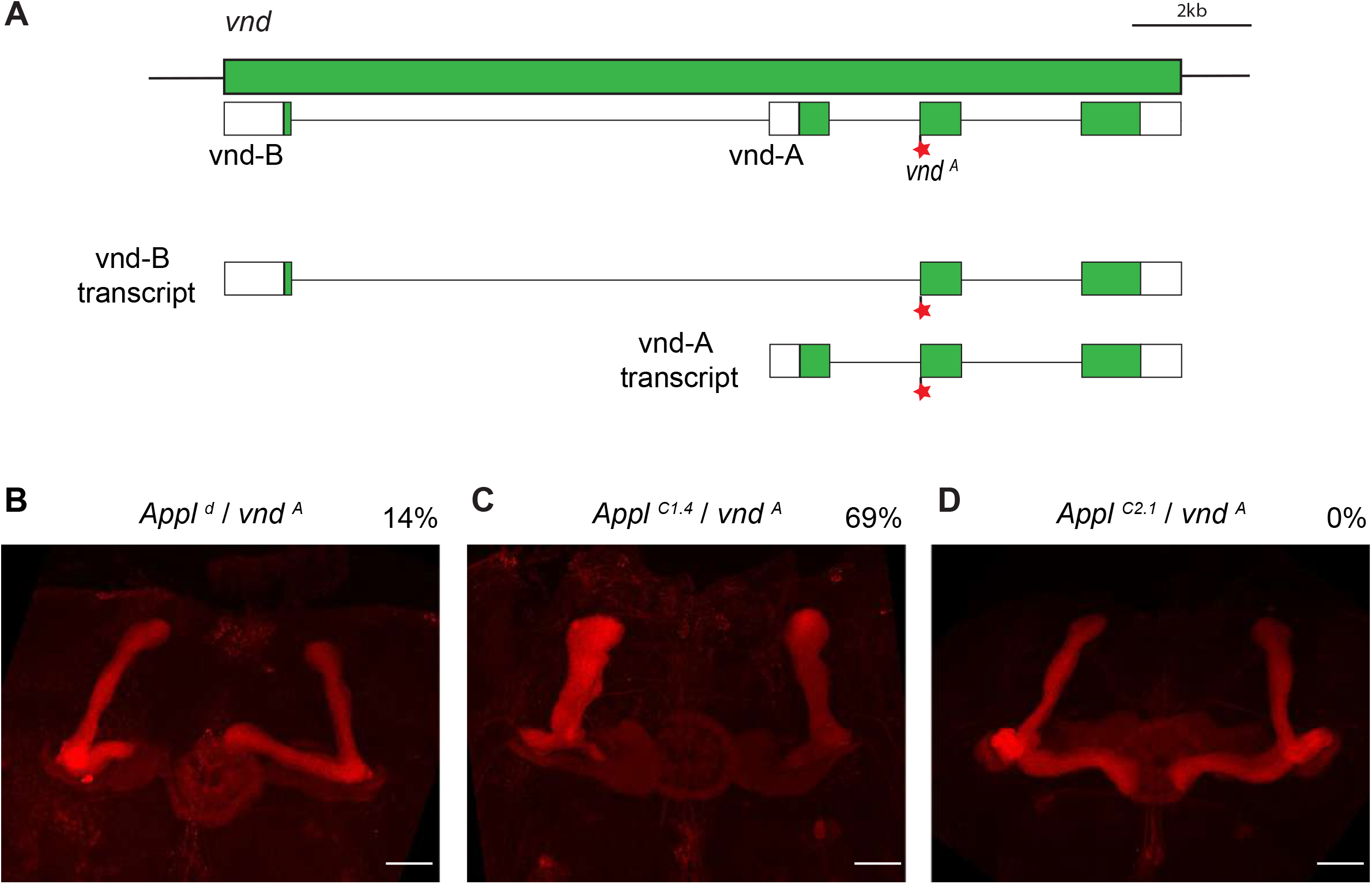
*Appl*^*d*^ and *Appl*^*C1*.*4*^, but not *Appl*^*C2*.*1*^, are impaired for *vnd* function. A) Schematic representation of *vnd* gene and transcripts. White boxes represent 5’UTR and 3’UTR, green boxes represent coding sequences. Red star represents point mutation on *vnd*^*A*^ allele. Schematic representation of *vnd-B* and *vnd-A* transcripts. The *vnd*^*A*^ mutation affects both transcripts. (B-D) Anti-Fas2 staining reveals the loss of β lobe on a representative *Appl*^*d*^/ *vnd*^*A*^ brain (n=214 MBs) (B), *Appl*^*C1*.*4*^/ *vnd*^*A*^ brain (n=147 MB) (C) but not on *Appl*^*C2*.*1*^/ *vnd*^*A*^ brain (n=248 MBs) (D). The % represents the proportion of loss of β lobe for each genotype. Scale bar = 50 µm. Full genotypes: (B) *Appl*^*d*^/y *vnd*^*A*^ *w* FRT19A*. (C) y *Appl*^*C1*.*4*^ *w*^*1118*^/ y *vnd*^*A*^ *w* FRT19A*. (D) *Appl*^*C2*.*1*^/ y *vnd*^*A*^ *w* FRT19A*.

### Generation of new *vnd* CRISPR alleles deleted either for *vnd-B* or *vnd-A* function

The two *vnd* transcripts, *vnd-A* and *vnd-B*, produce two different Vnd proteins, respectively Vnd-A and Vnd-B (Fig 2). It was proposed that, while Vnd-A has its main role during embryogenesis, Vnd-B acts during metamorphosis (Stepchenko et al., 2011). To determine which Vnd isoform is required for MB β-branch axon growth, we produced the new *vnd* alleles *vnd*^*CΔA*^ and *vnd*^*CΔB*^ by CRISPR/Cas9 genomic engineering that eliminate either the *vnd-A* or the *vnd-B* isoform, respectively (Fig 3 and S3 Fig). Males bearing a *vnd*^*CΔB*^ mutant allele (*vnd*^*CΔB*^/Y), as well as *vnd*^*CΔB*^/ *Appl*^*C1*.*4*^ females, are viable and have essentially wild-type MBs (Fig 3). Males bearing a *vnd*^*CΔA*^ mutant allele (*vnd*^*CΔA*^/Y) are embryonic lethal similarly to other *vnd* null alleles which affect both isoforms. *Appl*^*C1*.*4*^/*vnd*^*CΔA*^ female MBs displayed a strong phenotype of β lobe absence with a penetrance (64%) like those of *Appl*^*C1*.*4*^/*vnd*^*A*^ female MBs (Fig 4). We sought to determine the respective contribution of the lack of *Appl*^+^ and the lack of *vnd*^+^ for the strong β lobe absence in *Appl*^*C1*.*4*^/*vnd*^*CΔA*^ females. For this purpose, we added either a duplication of *Appl*^+^ or a duplication of *vnd*^+^ to this genotype. We found that *Appl*^*C1*.*4*^/*vnd*^*CΔA*^; *Dp-Appl*^+^/+ females displayed 69% of β lobe absence, a penetrance not distinguishable from that observed without the duplication of wildtype *Appl* (p = 0.42). Therefore, in these two types of females (*Appl*^*C1*.*4*^/*vnd*^*CΔA*^; +*/*+ *and Appl*^*C1*.*4*^/*vnd*^*CΔA*^; *Dp-Appl*^+^/+), the phenotype of β lobe absence seemed entirely due only to *vnd* lack-of-function. Indeed, *Appl*^*C1*.*4*^/*vnd*^*CΔA*^; +/+ females could be considered as *Appl*^*null*^ *vnd*^*strong*^/*Appl*^+^ *vnd*^*null*^ being homozygous mutant for *vnd* but heterozygous null for *Appl*. Similarly, *Appl*^*C1*.*4*^/*vnd*^*CΔA*^; *Dp-Appl*^+^/+ females could be considered as *Appl*^*null*^ *vnd*^*strong*^/*Appl*^+^ *vnd*^*null*^; *Dp-Appl*^+^/+ and are homozygous mutant for *vnd* but wild-type for *Appl*. Supporting the hypothesis of Vnd role in β axon growth, we observed a complete rescue when an extra dose of *vnd*^+^ is supplied as in *Appl*^*C1*.*4*^/*vnd*^*CΔA*^; *Dp-vnd*^+^/+ females (Fig 4). *Appl*^*C1*.*4*^/Y males have no *Appl*^+^ function (null allele) and display reduced *vnd*^+^ function. Moreover, *Appl*^*C1*.*4*^/Y male MB phenotype is moderately rescued by the presence of an extra dose of *Appl*^+^ but strongly rescued by an extra dose of *vnd*^+^ (Fig 4). Taken together, these results strongly indicate that the *vnd-A* transcript but not the *vnd-B* transcript is specifically required for MB β-branch axon growth.

**Fig 3.**
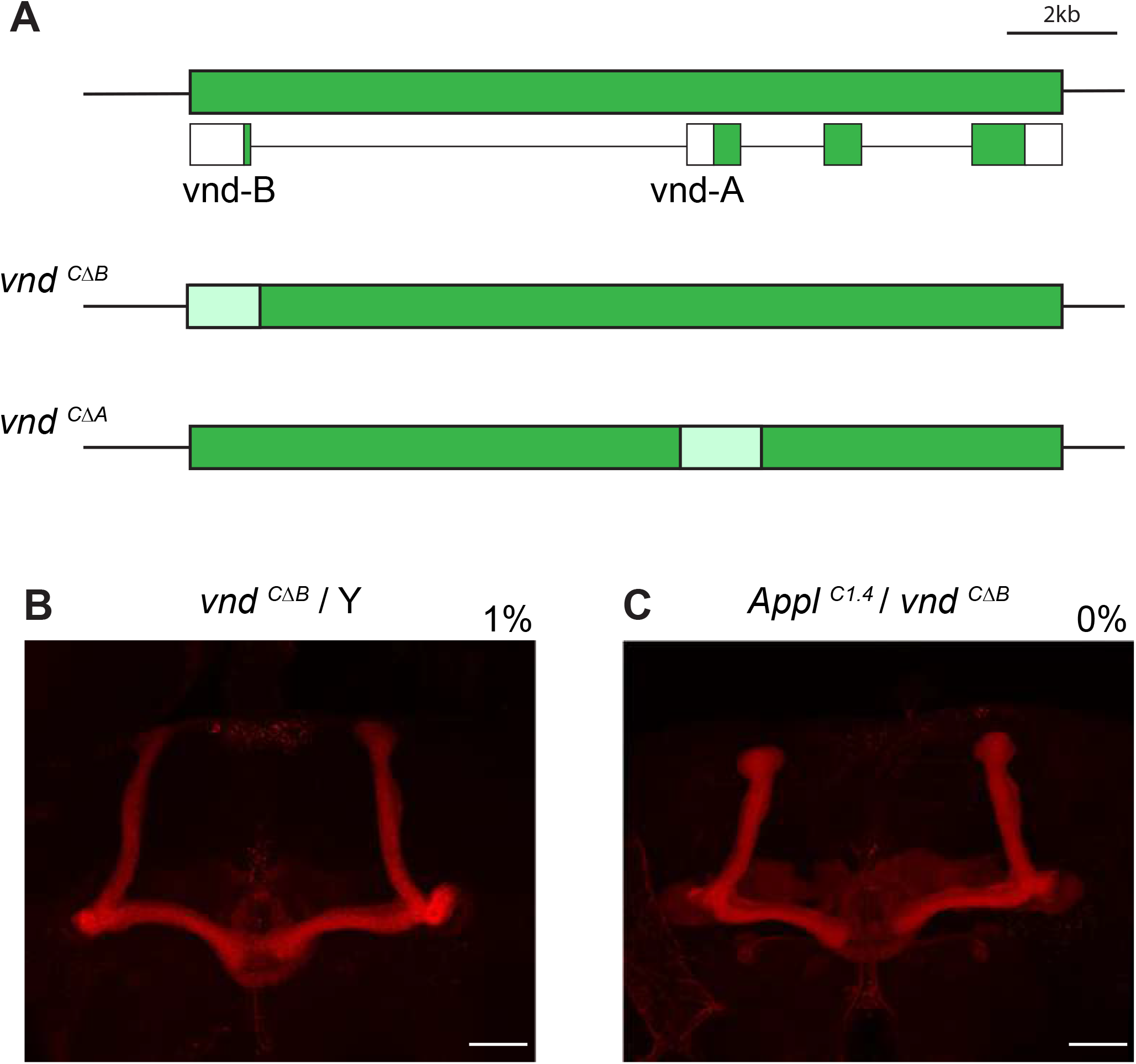
Generating new *vnd* CRISPR alleles deleted either for the *vnd-B* or the *vnd-A* function. (A) Schematic representation of *vnd* gene and transcripts. White boxes represent 5’UTR and 3’UTR, green boxes represent coding sequences. The deleted sequence region of *vnd*^*CΔA*^ and *vnd*^*CΔB*^ are represented in light green. (B-C) Anti-Fas2 staining reveals essentially wild-type looking MBs on a representative *vnd*^*CΔB*^ brain (n=146 MBs) (B) and *Appl*^*C1*.*4*^/ *vnd*^*CΔB*^ brain (n=159 MB) (C). The % represents the proportion of loss of β lobe for each genotype. Scale bar = 50 µm. Full genotypes: (B) *y vnd*^*CΔB*^ *w*^*1118*^/Y. (C) y *Appl*^*C1*.*4*^ *w*^*1118*^/ *y vnd*^*CΔB*^ *w*^*1118*^

**Fig 4.**
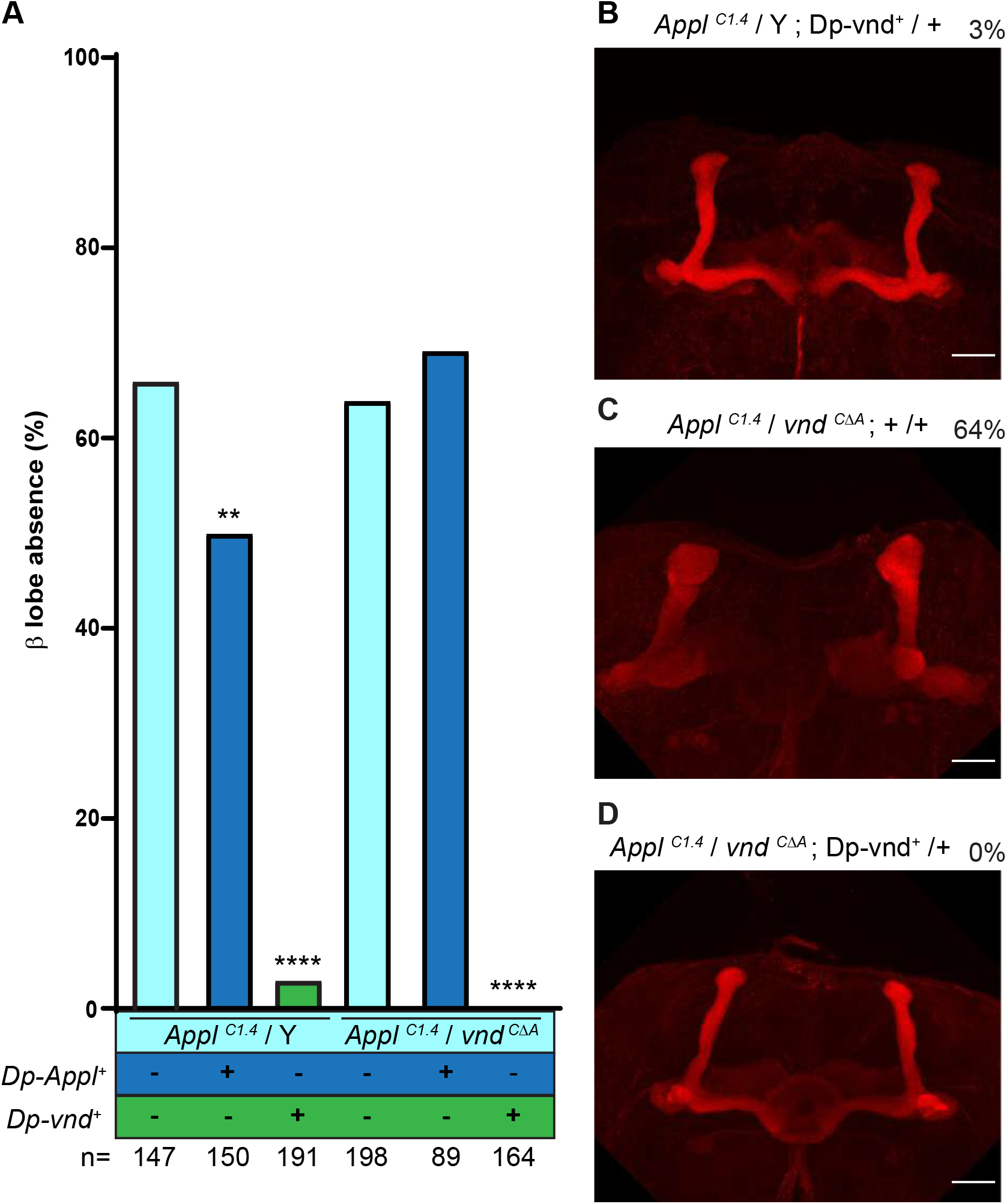
Vnd-A is required for MB β-branch axon outgrowth. (A) Quantitation of the rescue of MB β lobe absence of *Appl*^*C1*.*4*^/Y (66%; n= 147 MBs, see a representative brain in Fig.1D) and *Appl*^*C1*.*4*^/*vnd*^*CΔA*^ (64%; n=198 MBs) (light blue) by an *Appl* duplication (dark blue) (50%; n=150 MBs, 69%; n=89 MBs) and by a *vnd* duplication (green) (3%; n=191 MBs ; 0%; n=164 MBs). **p<0.01 and ****p<0.0001. (B-D) Anti-Fas2 staining on a representative rescued *Appl*^*C1*.*4*^/Y; *Dp-vnd*^*+*^/+ brain (B), *Appl*^*C1*.*4*^/*vnd*^*CΔA*^; +/+ brain (C) and on *Appl*^*C1*.*4*^/*vnd*^*CΔA*^ ; *Dp-vnd*^*+*^/+ brain (D). The % represents the proportion of loss of β lobe for each genotype. Scale bar = 50 µm. Full genotypes: (B) y *Appl*^*C1*.*4*^ *w*^*1118*^/Y; *Dp-vnd*^+^/+. (C) y *Appl*^*C1*.*4*^ *w*^*1118*^/*y vnd*^*CΔA*^ *w*^*1118*^; +/+. (D) y *Appl*^*C1*.*4*^ *w*^*1118*^/*y vnd*^*CΔA*^ *w*^*1118*^; *Dp-vnd*^+^/+.

### Vnd is expressed around the MBs and is not required within the MBs

In order to know where *vnd* is expressed, we employed a *vnd-T2A-GAL4* line where GAL4 is under the control of endogenous *vnd* regulatory sequences via CRISPR gene targeting so that GAL4 and *vnd* are translated from a single mRNA transcript (Chen et al., 2015; Lee et al., 2020). UAS-GFP labelling revealed a pattern in the ventral ganglion of the third larval instar central nervous system (L3 CNS) similar to that described with immunostaining with antibodies against Vnd (Stepchenko et al., 2011) thus validating the use of the *vnd-T2A-GAL4* line as a *bona fide vnd* reporter (S4 Fig). GFP was detected, in the developing brain, close to the MBs visualized by anti-Fas2 staining, from L3 to 24 h after puparium formation (APF) (Fig 5A, A’-5D, D’). Staining reveals a structure in the brain whose cell bodies are organized in a honeycomb pattern. To determine the identity of the cell bodies, we performed DAPI staining at the L3 stage. We observed a significant DAPI staining in these structures indicating that they correspond to cell nuclei (Fig 5E, E’). Moreover, these cell nuclei are Vnd positive and Repo negative (Fig 5F, F’-5G, G’) indicating that these cells correspond to neurons, rather than glia. The neurites emanating from these cell nuclei are very close to the developing MB medial lobe from which the adult β lobe develops (Fig 5). Noticeably, we did not observe any GFP labeling within the MBs themselves. This unexpected expression pattern suggests a non-cell-autonomous role for Vnd in the MB β-branch axon growth. We tested this hypothesis by MARCM mosaic analysis which allowed the generation of homozygous *vnd* loss-of-function MB clones in an otherwise heterozygous genetic background and overcame the homozygous lethality phenotype (Lee et al., 1999; Lee and Luo, 1999). Mitotic recombination was induced in late-stage embryos/early first instar larvae and the clones were analyzed at the adult stage. We obtained 20 MB *vnd*^*A/A*^ clones that include the αβ neurons. All the 20 *vnd* mutant clones displayed a β-branch axon growth that looked identical to that in wild-type clones (S5 Fig). Moreover, we obtained similar results, namely a normal β-branch axon growth, with two additional different null alleles, *vnd*^*Δ38*^ (8 clones) and *vnd*^*6*^ (4 clones) (S5 Fig). Taken together, these results demonstrate that, although *vnd* function is strongly required for MB β-branch axon growth, *vnd* is neither required nor expressed in the MBs themselves. Therefore, it is most likely that Vnd regulates MB axon growth by a non-cell-autonomous mechanism.

**Fig 5.**
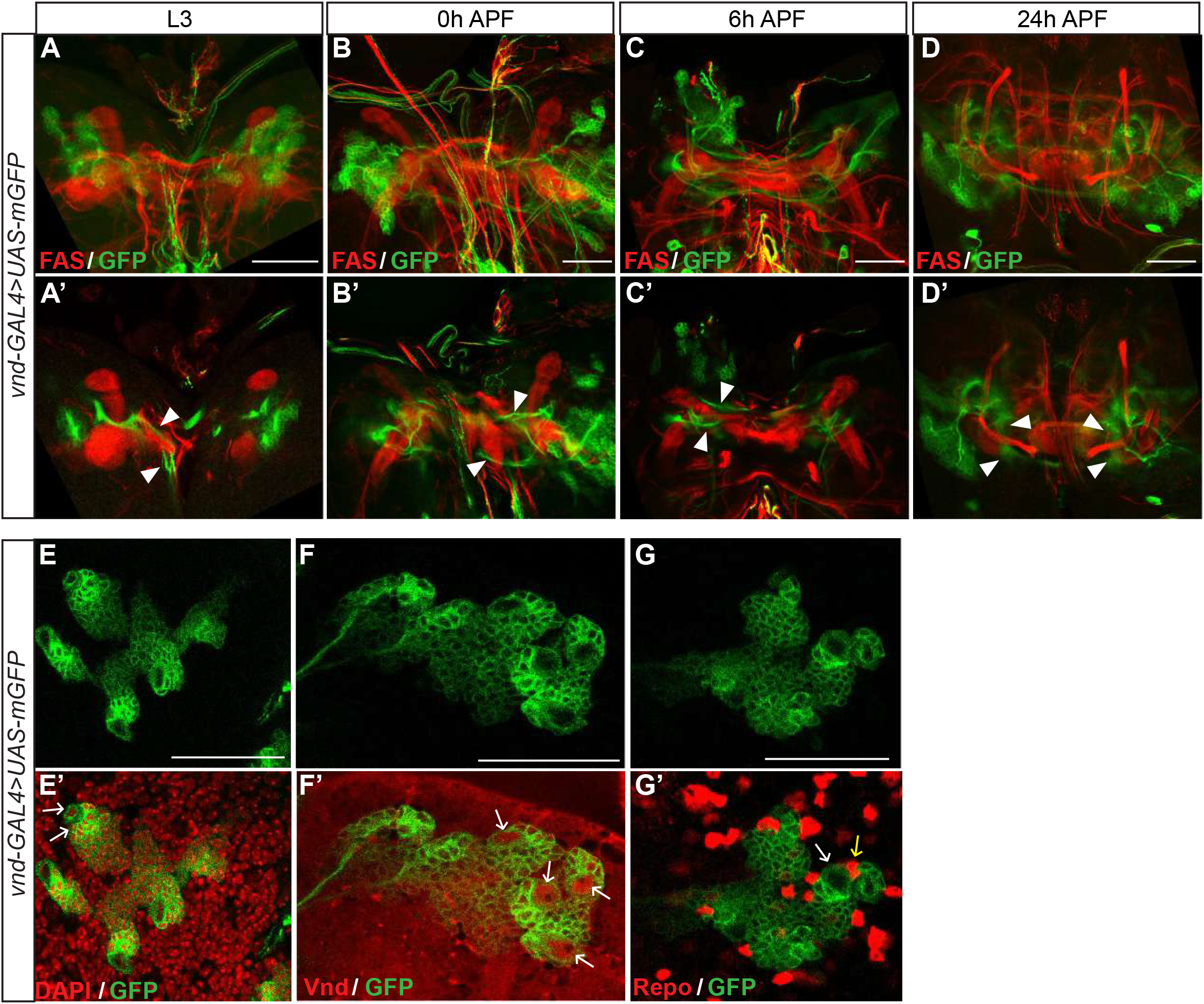
*vnd* is expressed close to the developing MB medial lobe. (A-D) Visualization of *vnd* expression in brain using *vnd-GAL4* driven *UAS-GFP* at L3, 0h, 6h, and 24h APF. GFP was visualized on green and anti-FasII staining labelling of MB axons on red. A-D are confocal z stacks taken the whole MB and A’-D’ are stacks of five sections from A-D respectively comprising GFP^+^ regions surrounding MB. Arrowheads point to *vnd-GAL4* driven *UAS-GFP* neurites that surround MB medial lobes, which correspond to γ lobes from L3 to 6 h APF (A’-C’) and β lobes at 24 h APF (D’). n ≥ 10 for each time point. (E-G) show single confocal sections showing *vnd* expression using *vnd-GAL4* driven *UAS-GFP* (single channel) at L3. E’ to G’ are merges of single channel E to G respectively and DAPI staining (red on E’), anti-vnd (red on F’) and anti-Repo (red on G’). DAPI labelled nuclei in GFP^+^ cell bodies are pointed by arrows in E’. These nuclei were also Vnd positive (arrows in F’) but were not labelled by an anti-Repo (white arrow in G’) which labels glia nuclei (red dots pointed by a yellow arrow in G’) indicating that these Vnd positive nuclei belong to neurons. Scale bars are 50 μm. n ≥ 5 brains for each type of staining. Full genotype: *vnd-T2A-GAL4 w\**; 2 x *UAS-mGFP*/+.

## Discussion

It has been proposed that Appl is part of the membrane complex formed by the core PCP proteins Fz and Vang (Soldano et al., 2013). *Appl*^*d*^ was, before this report, the only *Appl* null allele described. *Appl*^*d*^ mutant flies are completely lacking Appl function and exhibit 14.5 % MB β-lobe loss (Marquilly et al., 2021) in accordance with what has been reported (Liu et al., 2021; Soldano et al., 2013). This modestly penetrant axonal phenotype due to the lack of Appl could be due to partially redundant function provided by the other transmembrane receptors of the complex. Flies homozygous for the loss of function allele *Vang*^*stbm-6*^ exhibit 50% β-lobe loss (Liu et al., 2021). As the *Appl*^*d*^ allele involves complex chromosomic rearrangements (Luo et al., 1992), we leveraged CRISPR/Cas9 chromosomal engineering to produce a defined deletion of around 50 kb to specifically eliminate the *Appl* transcriptional unit. We recovered two *Appl* complete deletions (Fig 1): *Appl*^*C1*.*4*^ and *Appl*^*C2*.*1*^ removing around 52 kb and 47 kb and exhibiting 66 % and 9 % β-lobe loss respectively. DNA sequences analysis revealed that, while the *Appl*^*C2*.*1*^ deletion removes only *Appl* sequences, the *Appl*^*C1*.*4*^ deletion additionally removes a part of the proximally located transcriptional unit corresponding to the *vnd* gene (Fig 1). This indicates that in addition to Appl, *vnd* is likely involved in the β-branch axon outgrowth. Moreover, as can be expected from the extent of the deletions and based on genetic complementation tests, the *Appl*^*C1*.*4*^ deletion, but not the *Appl*^*C2*.*1*^ deletion, affects *vnd* function. Most importantly, the *Appl*^*d*^ deletion, which removes most of the Appl coding sequence and the intergenic region between *Appl* and *vnd*, also affects *vnd* function (Fig 1 and 2).

The role of *vnd* in β-branch axon outgrowth is complicated by the fact that the viable *vnd* mutant alleles (*Appl*^*d*^ *and* the *Appl*^*C1*.*4*^ deletion), do not encode functional Appl. Nevertheless, employing a small (less than 100 kb) genomic *Appl*^+^ duplication, we found that flies lacking only *vnd* function still exhibit 69% β-lobe loss and that this mutant phenotype is completely rescued by a small genomic *vnd*^+^ duplication (Fig 4). This demonstrates a requirement of *vnd* function in MB β-branch axon outgrowth. To better understand the roles of both Vnd isoforms in the MB β-branch axon outgrowth, we produced two mutations, *vnd*^*CΔB*^ and *vnd*^*CΔA*^, which remove the *vnd-B* and the *vnd-A* transcript respectively (Fig 3 and S3 Fig). *vnd-A* mRNA is expressed strongly in embryos and much less in larvae and adults contrarily to *vnd-B* mRNA (Stepchenko et al., 2011). It was therefore expected that the phenotype seen in adult MBs of pupal-born neurons should be due to *vnd-B* rather than *vnd-A*. Surprisingly, although Vnd-A has a significant role in MB axon growth, apparently Vnd-B plays very little to no role in this process. Noticeably, the MB phenotype of *Appl*^*C1*.*4*^/*vnd*^*CΔA*^ flies is similar in penetrance to that of *Appl*^*C1*.*4*^/*vnd*^*A*^ where *vnd*^*A*^ impacts both the *vnd-A* and *vnd-B* transcripts. This indicates that a lack of *vnd-A* function, and not that of vnd-B, underlies the poor β-branch axon outgrowth phenotype.

If Vnd interacts with Appl within the MBs, we hypothesized that it might be expressed within the MBs, and act cell-autonomously like Appl. However, endogenous *vnd* transcription monitored by GAL4 expression that is translated from the same mRNA transcript and was validated by anti-Vnd staining did not reveal Vnd expression within the MBs. Rather, *vnd* is transcribed in cells within a neuronal brain structure near the developing MB medial lobe from which the adult β lobe develops (Fig 5). Using MARCM mosaic analysis, we show that β axons extended from MB clones null for *vnd* function exhibit wild-type growth patterns (S5 Fig) demonstrating a non-cell-autonomous requirement for Vnd. Therefore, we favor the hypothesis that *vnd* expressed in a MB surrounding brain structure causes the expression of one or more secreted factors regulating the axon growth of the developing β medial lobe.

## Materials and methods

### Drosophila stocks

All crosses were performed on standard culture medium at 25^°^ C. Except where otherwise stated, all alleles and transgenes have been described previously (http://flystocks.bio.indiana.edu/). The following alleles were used: *Appl*^*C1*.*4*^, *Appl*^*C2*.*1*^, *vnd*^*CΔA*^ and *vnd*^*CΔB*^ were generated in this study. *Appl-y*^+^ (BL #56073), *Appl*^*d*^ (BL #43632), *vnd*^*A*^ (BL #57139), *vnd*^*Δ38*^ (Chu et al., 1998), *vnd*^*6*^ (Jimenez et al., 1995). The following transgenes were used: *UAS-mCD8::GFP* (BL #5137) and the duplications resulting from genomic transgenes *Dp(1;3) DC430* (BL #32288) and *Dp(1*.*3) DC009* (BL #30219) which are both inserted into the same attP site at 65B2. For simplicity, we refer to *Dp(1;3) DC430* as *Dp-Appl*^+^ and *Dp(1*.*3) DC009* as *Dp-vnd*^+^. *Dp-Appl*^+^ is ∼ 97 kb and contains 11 genes including the *Appl* gene but not the *vnd* gene. *Dp-vnd*^+^ is also ∼ 97 kb and contains 15 genes including the *vnd* gene but not the *Appl* gene. We used two GAL4 lines: *vnd-T2A-GAL4* (Lee et al., 2020) which reveals *vnd* transcription and *c739-GAL4* (BL #7362) expressed in adult αβ MB neurons (Aso et al., 2009). Recombinant chromosomes were obtained by standard genetic procedures. In the case of the recombination between *y*^+^ and *ApplC*^*2*.*1*^ (∼ 170 kb apart) we screened for [*y*^+^] viable males coming from *y*^+^ *vnd*^*Δ38*^/*y ApplC*^*2*.*1*^ females. Note that *vnd*^*Δ38*^ is an embryonic lethal mutation. Four [*y*^+^] viable males out of 11320 males were obtained. One male was sterile, but three *y*^+^ *ApplC*^*2*.*1*^ /Y males gave progeny (percentage of recombination circa 0.027%) and were confirmed by genomic PCR. MARCM stocks: *tubP-GAL80, hsFLP122, hsFLP1, FRT19A* (all on the X chromosome).

### CRISPR-Cas9 strategy

All guide RNA sequences (single guide RNA (sgRNA)) were selected using the algorithm targetfinder.flycrispr.neuro.brown.edu/ containing 20 nucleotides each (PAM excluded) and are predicted to have zero off-targets. We selected one pair of sgRNA for the *Appl* gene and two pairs of sgRNA for the *vnd* gene. For *Appl*, the pair is targeting the entire transcriptional unit (S2 Fig). For *vnd*, each pair is targeting either the A specific region of *vnd* or the B specific region of *vnd* (S3 Fig). We used the following oligonucleotide pairs: CRISPR-1 Appl fwd and CRISPR-1 Appl rev to target the all *Appl* region, CRISPR-1 vnd A fwd and CRISPR-1 vnd A rev to target the A region of *vnd*, CRISPR-1 vnd B fwd and CRISPR-1 vnd B rev to target the B region of *vnd* (see the corresponding oligonucleotide sequences in S2 and S3 Fig.). We introduced two sgRNA sequences into pCFD4 (Port et al., 2014), a gift from Simon Bullock (Addgene plasmid # 49411) by Gibson Assembly (New England Biolabs) following the detailed protocol at crisprflydesign.org. For PCR amplification, we used the protocol described on that website. Construct injection was performed by Bestgene (Chino Hills, CA) and all the transgenes were inserted into the same attP site (VK00027 at 89E11). Being aware that the DNA excision of nearly 50 kb might be rare, we have decided to use a positive screen before validation by genomic PCR. We have used a stock where the *Appl* gene is marked by a *y*^+^ construct. The deletion of the *Appl* gene should therefore be associated to *y*^*-*^ phenotype. Transgenic males expressing the *Appl* sgRNAs and bearing an isogenized X chromosome (*y Appl-y*^*+*^ *Act-Cas9 w\**) were crossed to *FM7c/ph*^*0*^ *w* females. 180 crosses were set up and only two crosses gave rise to *y Appl*^*deletion?*^ *Act-Cas9 w*/FM7c* females indicating 1% of CRISPR efficacy. From each of these two crosses, a single *y Appl*^*deletion*^ *Act-Cas9 w* /FM7c* female was crossed with *FM7c* males to make a stock (*Appl*^*C1*.*4*^ and *Appl*^*C2*.*1*^ where Cas9 was removed). The deletion was then validated by genomic PCR using two pairs of primers: *Appl*^*C1*.*4*^ fwd (GAGCCAGATACACAAGCACA) / *Appl*^*C1*.*4*^ rev (GGCTTTGTTTACTTCCTGGC) and *Appl*^*C2*.*1*^ fwd (TCCTACTACGTTCCACAATC) / *Appl*^*C2*.*1*^ rev (TAATGCCCAACATATCCAAC). The precise endpoints of the deletion were determined by sequencing (Genewiz, France). Transgenic males expressing the different *vnd* sgRNAs were crossed to *y nos-Cas9 w\** females bearing an isogenized X chromosome. 100 crosses were set up for each sgRNA pair, with up to 5 males (because of poor viability) containing both the sgRNAs and *nos-Cas9*, and 5 *FM7c/ph*^*0*^ *w* females. From each selected cross, a single *y vnd*^*deletion?*^ *nos-Cas9 w* /FM7c* female was crossed with *FM7c* males to make a stock which was validated for the presence of an indel by genomic PCR with primers flanking the anticipated deletion and the precise endpoints of the deletion were determined by sequencing (Genewiz, France) using *vnd*-specific primers: *vnd*^*CΔA*^ fwd (ccaacaaagccgagagtctct) / *vnd*^*CΔA*^ rev (cgggaatttctaagccagggt) and *vnd*^*CΔB*^ fwd (cgatttggggcgttgtgagta) / *vnd*^*CΔB*^ rev (gttgggctttaatccgggagt). A CRISPR efficacy of at least 5% was obtained in both cases. We kept a single stock from each set of CRISPR experiment: *vnd*^*CΔA*^ and *vnd*^*CΔB*^ where Cas9 was removed.

### Brain dissection, immunostaining and MARCM mosaic analysis

Adult fly heads and thoraxes were fixed for 1 h in 3.7% formaldehyde in phosphate-buffered saline (PBS) and brains were dissected in PBS. For larval and pupal brains, brains were first dissected in PBS and then fixed for 15 min in 3.7% formaldehyde in PBS. They were then treated for immunostaining as described (Boulanger et al., 2011; Lee and Luo, 1999). Antibodies, obtained from the Developmental Studies Hybridoma Bank, were used at the following dilutions: Mouse monoclonal anti–Fas2 (1D4) 1:10, mouse monoclonal anti-Repo (8D1.2) 1:10 and guinea pig polyclonal anti-vnd (1/1000). Goat secondary antibodies conjugated to Cy3 against mouse or guinea pig IgG (Jackson Immunoresearch laboratory) were used at 1:300 for detection. DAPI (Sigma) was used after secondary antibody washes. Tissues were incubated for 10 minutes at room temperature in a 1/1000 solution from a stock containing 1 mg/ml of DAPI solution, then washed. To generate clones in the MB, we used the MARCM technique (Lee and Luo, 1999). First instar larvae were heat-shocked at 37°C for 1 h. Adult brains were fixed for 15 min in 3,7% formaldehyde in PBS before dissection and GFP visualization.

### Production of the anti-Vnd antiserum

Full length vnd-RA was amplified via PCR from cDNA and cloned into a pRSET bacterial expression vector (ThermoFisher) in frame with an N-terminal 6x-His tag, amplified in DH5alpha and transformed into BL21 competent cells for protein production. His-tagged VndA was purified over a Nickel-NTA agarose resin (Qiagen) according to manufacturer’s recommendation. Purified protein was injected into guinea pigs by Squarix GmbH (Marl, Germany, www.squarix.de), selected by determining that pre-immunization sera did not produce signal in Western blots of fly extracts. Successive bleeds were screened for distinct signal in Western blots using embryo extracts and by immunohistochemistry, where signal was expected post 2.5hrs AED.

### Microscopy and image processing

Images were acquired at room temperature using a Zeiss LSM 780) equipped with a 40x PLAN apochromatic 1.3 oil-immersion differential interference contrast objective lens and a Leica SP8 laser scanning confocal microscopes (MRI Platform, Institute of Human Genetics, Montpellier, France. The immersion medium used was Immersol 518F. The acquisition software used was Zen 2011 (black edition) for the Zeiss and LasX for the Leica. Contrast and relative intensities of the green (GFP), of the red (Cy3) and of the DAPI channels were processed with ImageJ (Fiji) software. Settings were optimized for detection without saturating the signal.

### Statistics

Comparison between two groups expressing a qualitative variable was analyzed for statistical significance using the Chi^2^ test. (BiostaTGV: http://biostatgv.sentiweb.fr/?module=tests). Values of p < 0.05 were considered to be significant. Statistical significance was defined as: ns = not statistically different, **p<0.01 and****p<0.0001.

## Acknowledgments

We thank Aymeric Chartier (IGH, Montpellier) for fruitful discussions regarding the generation of the *Appl* CRISPR alleles. We thank the Bloomington *Drosophila* Stock Center (Indiana University), the BioCampus RAM-*Drosophila* facility and the BioCampus imaging facility MRI (Montpellier, France). We acknowledge BestGene and Genewiz for transgene service and DNA sequencing respectively. The 1D4 anti-Fasciclin II hybridoma and the 8D12 anti-Repo monoclonal antibody developed by Corey Goodman were obtained from the Developmental Studies Hybridoma Bank, created by the NICHD of the NIH and maintained at the University of Iowa, Department of Biology, Iowa City, IA 52242. We thank Chris Doe for the *vnd*^*Δ38*^ and *vnd*^*6*^ alleles and Tzumin Lee for the *vnd-T2A-GAL4* line.

## Supporting information

**S1 Fig.**
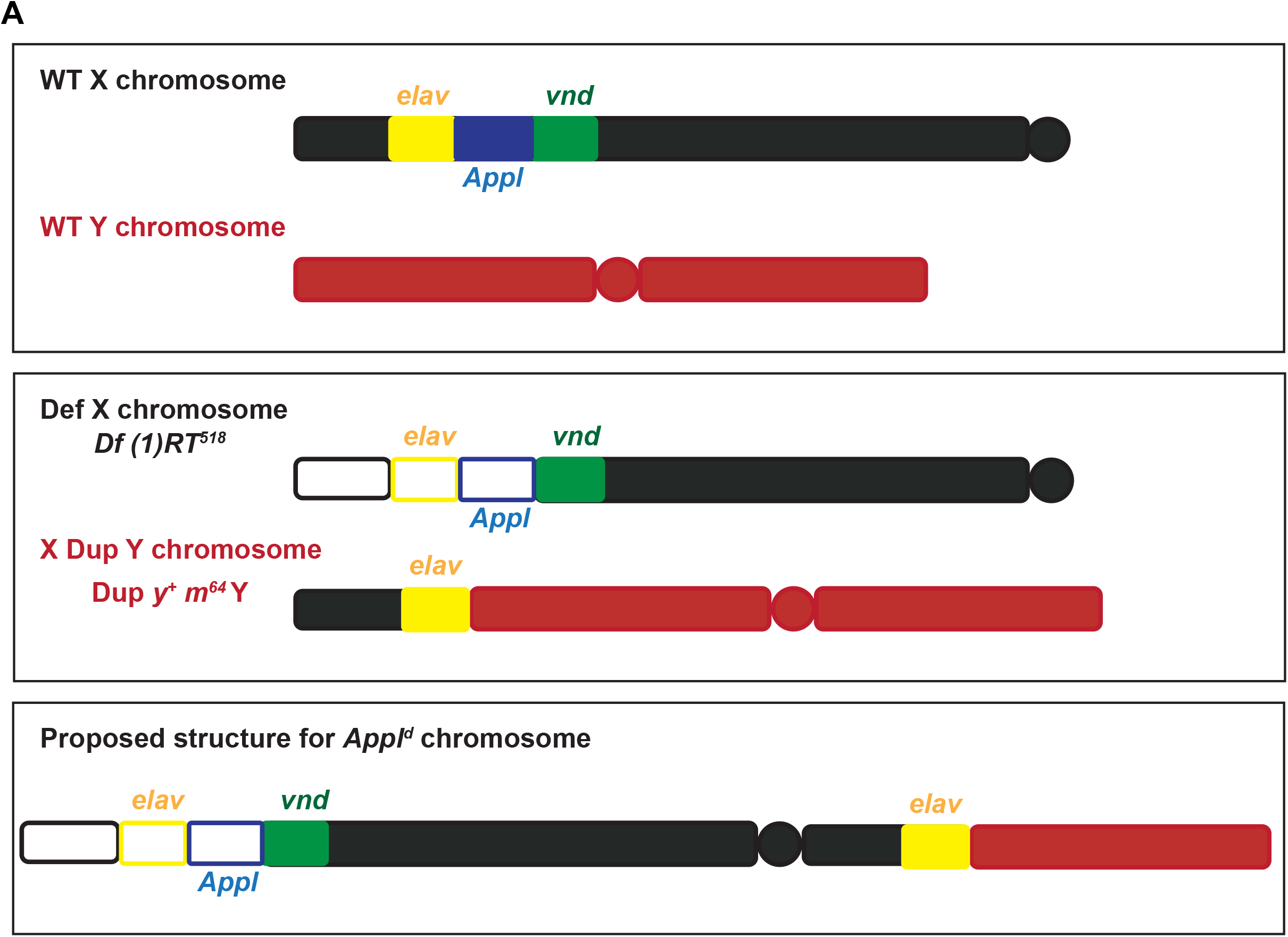
Scheme of *Appl*^*d*^ chromosomic organization. (A) Schematic representation of *Appl*^*d*^ mutant allele. The upper panel represents the wild-type (WT) X (black) and Y (orange) chromosomes. The *elav, Appl* and *vnd* genes are represented respectively in yellow, blue, and green boxes. The middle panel represents the deficiency Df (1) RT518 on the X chromosome deleting the distal part of the chromosome including the *elav* and *Appl* genes. The X Dup Y chromosome represents a duplication of the X chromosome linked to the Y chromosome including the distal part of the X chromosome including the *elav* gene. The lower panel represents a proposed structure of the *Appl*^*d*^ chromosome where the deficiency-bearing X chromosome had been linked by gamma irradiation to the X Dup Y chromosome (adapted from (Luo et al., 1992)). Additionally, we found that the Y chromosome behaves genetically as linked to the X chromosome.

**S2 Fig.**
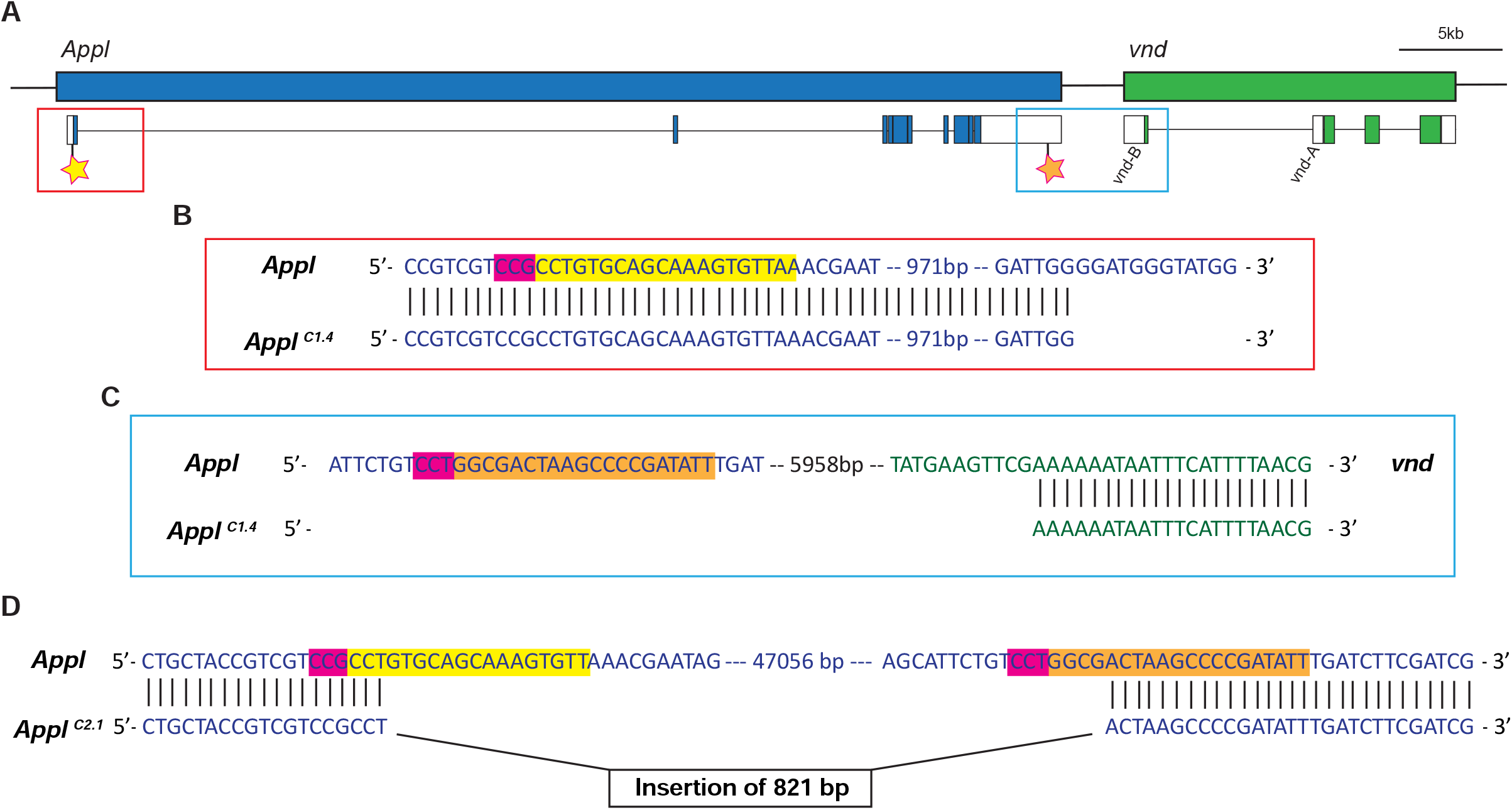
Structure of the *Appl*^*C1*.*4*^ and *Appl*^*C2*.*1*^ CRISPR alleles. (A) Schematic representation of *Appl* (blue) and *vnd* (green) genes and transcripts. White boxes represent 5’UTR and 3’UTR, blue and green boxes represent *Appl* and *vnd* coding sequences respectively. The yellow stars in *Appl* 5’ region and the orange star in *Appl* 3’ region represent the position of the sgRNA used to create *Appl*^*C1*.*4*^ and *Appl*^*C2*.*1*^ mutants. The red and blue boxes represent respectively the regions magnified in B and C. (B) Representation of the 5’ sequence targeted by the sgRNA (yellow) and the PAM domain (magenta) in the WT chromosome *Appl* (blue) (upper line). The lower line is the sequence of the *Appl*^*C1*.*4*^ mutant showing the beginning of the deletion about 1kb after the region targeted by the sgRNA. (C) Representation of the 3’ sequence targeted by the sgRNA (orange) and the PAM domain (magenta) in the WT chromosome *Appl* (blue) and *vnd* (green) (upper line). The lower line is the sequence of the *Appl*^*C1*.*4*^ mutant showing the end of the deletion about 6kb after the region targeted by the sgRNA. (D) Representation of the 5’ and 3’ sequences targeted by the sgRNA (yellow in 5’ and orange in 3’) and the PAM domain (magenta) in the WT chromosome *Appl* (blue) (upper line). The lower line is the sequence of the *Appl*^*C2*.*1*^ mutant showing the beginning of the deletion in the sequence targeted by the 5’ sgRNA (yellow) and the end of the deletion in the sequence targeted by the 3’ sgRNA (orange). Circa 46 kb of *Appl* transcribed sequence including all the coding sequence has been entirely deleted and replaced by a sequence of 821bp (see below) primarily derived from the fly retrotransposon micropia-like LTR polyprotein gene (NCBI Blast). ATAAACGCGTCATTAAGCTGTTCCATTTTCTAATCGCGCAGCGAGTTTCGGTGGTCGAAGTCTAGAATTGAAAAAGAAGAAGAAGAAGATACTCTGTATGTTATGGTTAAGAATAAAAGCTTAAGTTCAGTTTGTGTTATGAAACACAAAATGTGCCTTTGAACTTGATATTGGTCATTACACGAGGATGTGTAAATGTCAGATGGTTATCCGTCTATGAGAGACGATGGTTATCCGTCTATGAGAGACGATGGTTATCCGTCTATGAGAGACGATGGTTATCCGTCCATGAGAGACGATGGTTATCCGTCTATGAGAGACGATGGTTATCCGTCTATGAGAGACGATGGTTATCCACCTAAGACAGACGATGATTTCAGTTTAGAATCTCTTGTTACGTAAGAAATTTGAATCAAAATTGGGAGTACTACACGAGGACGTGTCAAGGTCAGGATGGCCGTGTCGGAGCATACTGTATGTACGCTCTGTATGTGATTGTTGCTGCGTATTGCACGTACGCGCTGTGTGTGTATTGCTGTGCATGGGAAATCTGGTTTTTACCAGAATTCCTAATTTCATCATATTAGTAATATTGCATCAGCCGTTAATTGCGCTCTGATTGGTCCTTTGCATTTTGTCGCTTTAAGAAGTACCGTTTTAGCGAGACGAAGCGCTACAACGCTGTCGTCGTGTGTGGTTCGGGAGCTATGCGGGGAAGTAAGTCTGGCGTTGTCACGCAGTTTTTATAGAAACGTG AAAGCATAAGGCACCGAAACTCGCTTCGCTAGGTACGCATATGGCACAAAATAATCAAGTTCTA

**S3 Fig.**
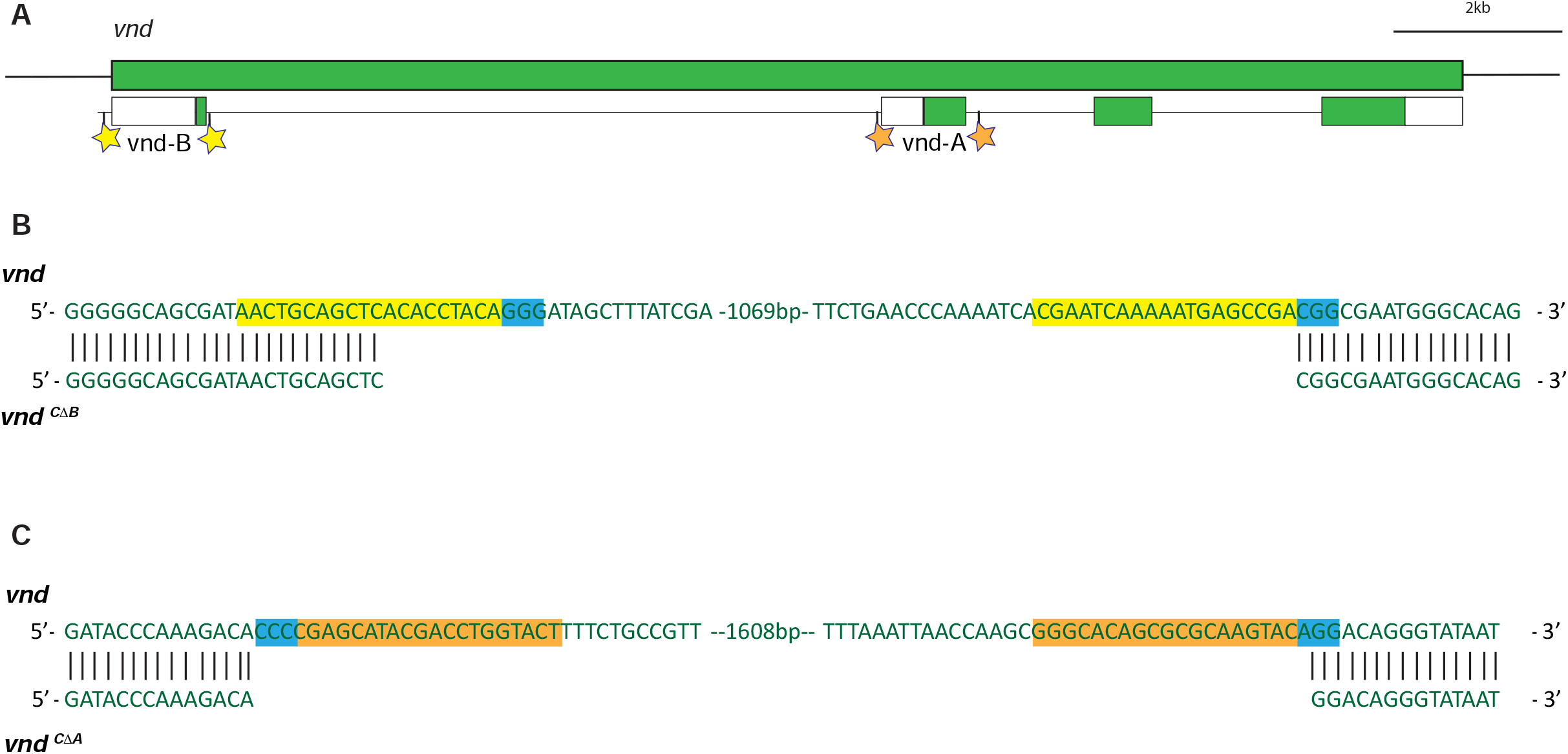
Structure of the *vnd*^*CΔB*^ and *vnd*^*CΔA*^ CRISPR alleles. (A) Schematic representation of *vnd* (green) gene and transcripts. White boxes represent 5’UTR and 3’UTR, green boxes represent *vnd* coding sequences. The yellow stars represent the position of the sgRNA used to create *vnd*^*CΔB*^ mutant. The orange stars represent the position of the sgRNA used to create *vnd*^*CΔA*^ mutant. (B) Representation of the sequences targeted by the sgRNA (yellow) and the PAM domain (blue) in the WT chromosome *vnd* (green) (upper line). The lower line is the sequence of the *vnd*^*CΔB*^ mutant showing the beginning of the deletion in the sequence targeted by the 5’ sgRNA (yellow) and the end of the deletion in the sequence targeted by the 3’ sgRNA (yellow). (C) Representation of the sequences targeted by the sgRNA (orange) and the PAM domain (blue) in the WT chromosome *vnd* (green) (upper line). The lower line is the sequence of the *vnd*^*CΔA*^ mutant showing the beginning of the deletion at one nucleotide from the sequence targeted by the 5’ sgRNA (orange) and the end of the deletion within in the sequence targeted by the 3’ sgRNA (orange).

**S4 Fig.**
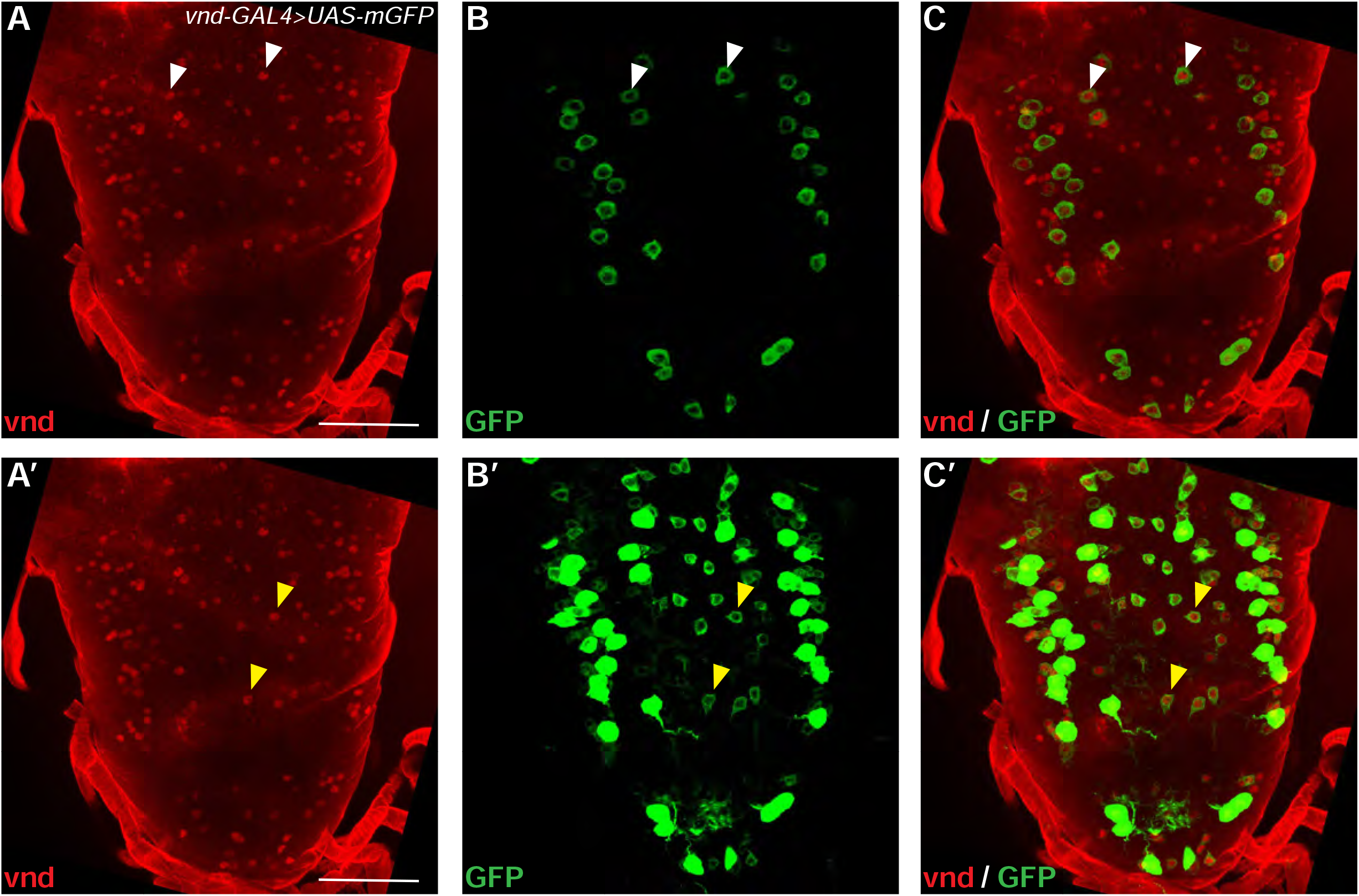
Vnd L3 CNS labeling. (A-C) Visualization of *vnd* expression in VNC using both *vnd-GAL4* driven *UAS-GFP* (green) and an anti-vnd antibody (red) at L3 stages. Images A-C and A’-C’ are the same confocal z-projection. Laser intensities leading to the green color were increased from B, C to B’, C’ to illustrate the high degree of colocalization between the two markers. Low laser intensity allowed us to show red nuclear vnd staining in only some GFP^+^ cell bodies (white arrowheads in A-C). Soma containing vnd^+^ positive nuclei that do not seem to be surrounded by GFP in A-C, can be visualized by increased green laser intensity A’-C’ (yellow arrowheads), which saturates somas visualized in A-C. Scale bars are 50 μm. n ≥ 5 brains. Full genotype: *vnd-T2A-GAL4 w\**; 2 x *UAS-mGFP*/+.

**S5 Fig.**
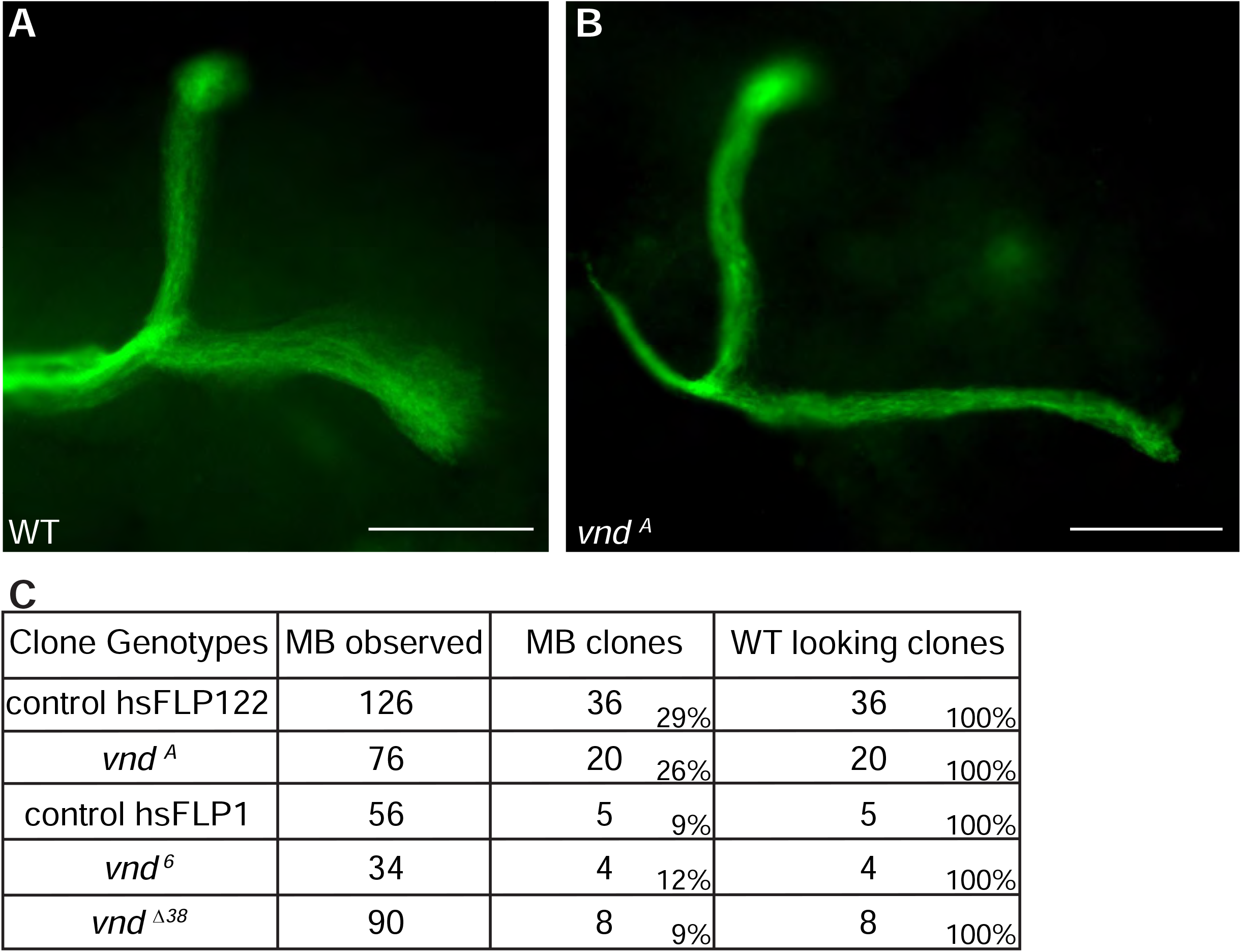
*vnd* null mutant clones. (A-B) A representative neuroblast MARCM clone in a wild-type (WT) brain (A) and in a *vnd*^*A*^ brain (B). Scale bar = 50 µm. (C) Number of clones obtained in WT, *vnd*^*A*^, *vnd*^*6*^ and *vnd*^*Δ38*^. Full genotypes: (A) *w* tubP-GAL80 hsFLP122 FRT19A/w*^*1118*^ *sn FRT19A; c739-GAL4 UAS-mCD8-GFP/*+ (also control hsFLP122 in C). (B) *w* tubP-GAL80 hsFLP122 FRT19A/y vnd*^*A*^ *w FRT19A; c739-GAL4 UAS-mCD8-GFP/*+ (also vnd^A^ in C). (C) control hsFLP1: *w tubP-GAL80 hsFLP1 FRT19A* (from the Bloomington stock 5133)/*w UAS-mCD8-GFP sn FRT19A*; *c739-GAL4 UAS-mCD8-GFP/*+. vnd^6^: *w tubP-GAL80 hsFLP1 FRT19A/y vnd*^*6*^ *w UAS-mCD8-GFP sn FRT19A*; *c739-GAL4 UAS-mCD8-GFP/*+. vnd^Δ38^: *w tubP-GAL80 hsFLP1 FRT19A/vnd*^*Δ38*^ *w UAS-mCD8-GFP FRT19A*; *c739-GAL4 UAS-mCD8-GFP/*+.

## References

Aso, Y., Grubel, K., Busch, S., Friedrich, A. B., Siwanowicz, I. and Tanimoto, H. (2009). The mushroom body of adult Drosophila characterized by GAL4 drivers. J Neurogenet 23, 156–172.

Boulanger, A., Clouet-Redt, C., Farge, M., Flandre, A., Guignard, T., Fernando, C., Juge, F. and Dura, J. M. (2011). ftz-f1 and Hr39 opposing roles on EcR expression during Drosophila mushroom body neuron remodeling. Nat Neurosci 14, 37–44.

Cassar, M. and Kretzschmar, D. (2016). Analysis of Amyloid Precursor Protein Function in Drosophila melanogaster. Front Mol Neurosci 9, 61.

Chen, H. M., Huang, Y., Pfeiffer, B. D., Yao, X. and Lee, T. (2015). An enhanced gene targeting toolkit for Drosophila: Golic+. Genetics 199, 683–694.

Chu, H., Parras, C., White, K. and Jimenez, F. (1998). Formation and specification of ventral neuroblasts is controlled by vnd in Drosophila neurogenesis. Genes Dev 12, 3613–3624.

Doudna, J. A. and Charpentier, E. (2014). Genome editing. The new frontier of genome engineering with CRISPR-Cas9. Science 346, 1258096.

Haelterman, N. A., Jiang, L., Li, Y., Bayat, V., Sandoval, H., Ugur, B., Tan, K. L., Zhang, K., Bei, D., Xiong, B., et al. (2014). Large-scale identification of chemically induced mutations in Drosophila melanogaster. Genome Res 24, 1707–1718.

Jimenez, F., Martin-Morris, L. E., Velasco, L., Chu, H., Sierra, J., Rosen, D. R. and White, K. (1995). vnd, a gene required for early neurogenesis of Drosophila, encodes a homeodomain protein. EMBO J 14, 3487–3495.

Lee, T., Lee, A. and Luo, L. (1999). Development of the Drosophila mushroom bodies: sequential generation of three distinct types of neurons from a neuroblast. Development 126, 4065–4076.

Lee, T. and Luo, L. (1999). Mosaic analysis with a repressible cell marker for studies of gene function in neuronal morphogenesis. Neuron 22, 451–461.

Lee, Y. J., Yang, C. P., Miyares, R. L., Huang, Y. F., He, Y., Ren, Q., Chen, H. M., Kawase, T., Ito, M., Otsuna, H., et al. (2020). Conservation and divergence of related neuronal lineages in the Drosophila central brain. Elife 9.

Leyssen, M., Ayaz, D., Hebert, S. S., Reeve, S., De Strooper, B. and Hassan, B. A. (2005). Amyloid precursor protein promotes post-developmental neurite arborization in the Drosophila brain. EMBO J 24, 2944–2955.

Lin, S. (2023). The making of the Drosophila mushroom body. Front Physiol 14, 1091248.

Liu, T., Zhang, T., Nicolas, M., Boussicault, L., Rice, H., Soldano, A., Claeys, A., Petrova, I., Fradkin, L., De Strooper, B., et al. (2021). The amyloid precursor protein is a conserved Wnt receptor. Elife 10.

Luo, L., Tully, T. and White, K. (1992). Human amyloid precursor protein ameliorates behavioral deficit of flies deleted for Appl gene. Neuron 9, 595–605.

Marquilly, C., Busto, G. U., Leger, B. S., Boulanger, A., Giniger, E., Walker, J. A., Fradkin, L. G. and Dura, J. M. (2021). Htt is a repressor of Abl activity required for APP-induced axonal growth. PLoS Genet 17, e1009287.

Nicolas, M. and Hassan, B. A. (2014). Amyloid precursor protein and neural development. Development 141, 2543–2548.

Port, F., Chen, H. M., Lee, T. and Bullock, S. L. (2014). Optimized CRISPR/Cas tools for efficient germline and somatic genome engineering in Drosophila. Proc Natl Acad Sci U S A 111, E2967–2976.

Rabah, Y., Berwick, J. P., Sagar, N., Pasquer, L., Placais, P. Y. and Preat, T. (2025). Astrocyte-to-neuron H(2)O(2) signalling supports long-term memory formation in Drosophila and is impaired in an Alzheimer’s disease model. Nat Metab 7, 321–335.

Selkoe, D. J. and Hardy, J. (2016). The amyloid hypothesis of Alzheimer’s disease at 25 years. EMBO Mol Med 8, 595–608.

Soldano, A. and Hassan, B. A. (2014). Beyond pathology: APP, brain development and Alzheimer’s disease. Curr Opin Neurobiol 27, 61–67.

Soldano, A., Okray, Z., Janovska, P., Tmejova, K., Reynaud, E., Claeys, A., Yan, J., Atak, Z. K., De Strooper, B., Dura, J. M., et al. (2013). The Drosophila Homologue of the Amyloid Precursor Protein Is a Conserved Modulator of Wnt PCP Signaling. PLoS biology 11, e1001562.

Stepchenko, A. G., Pankratova, E. V., Doronin, S. A., Gulag, P. V. and Georgieva, S. G. (2011). The alternative protein isoform NK2B, encoded by the vnd/NK-2 proneural gene, directly activates transcription and is expressed following the start of cells differentiation. Nucleic Acids Res 39, 5401–5411.

